# Dopamine and acetylcholine correlations in the nucleus accumbens depend on behavioral task states

**DOI:** 10.1101/2024.05.03.592439

**Authors:** Kauê Machado Costa, Zhewei Zhang, Yizhou Zhuo, Guochuan Li, Yulong Li, Geoffrey Schoenbaum

**Affiliations:** National Institute on Drug Abuse Intramural Research Program, National Institutes of Health, Baltimore, MD, 21224, USA; State Key Laboratory of Membrane Biology, School of Life Sciences, Peking University, Beijing 100871, China; PKU-IDG/McGovern Institute for Brain Research, Beijing, 100871, China; Peking-Tsinghua Center for Life Sciences, New Cornerstone Science Laboratory, Academy for Advanced Interdisciplinary Studies, Peking University Beijing, 100871, China; Chinese Institute for Brain Research, Beijing, 102206, China; Institute of Molecular Physiology, Shenzhen Bay Laboratory, Shenzhen, Guangdong, 518055, China; National Biomedical Imaging Center, Peking University, Beijing, 100871, China

**Keywords:** Dopamine, acetylcholine, rat, fiber photometry, instrumental, approach, ramps, reward prediction error, salience, motivation

## Abstract

Dopamine in the nucleus accumbens ramps up as animals approach desired goals. These ramps have received intense scrutiny because they seem to violate long-held hypotheses on dopamine function. Furthermore, it has been proposed that they are driven by local acetylcholine release, i.e., that they are mechanistically separate from dopamine signals related to reward prediction errors. Here, we tested this hypothesis by simultaneously recording accumbal dopamine and acetylcholine signals in rats executing a task involving motivated approach. Contrary to recent reports, we found that dopamine ramps were not coincidental with changes in acetylcholine. Instead, we found that acetylcholine could be positively, negatively, or uncorrelated with dopamine depending on whether the task phase was determined by a salient cue, reward prediction error, or active approach, respectively. Our results suggest that accumbal dopamine and acetylcholine are largely independent but may combine to engage different postsynaptic mechanisms depending on the behavioral task states.

## Introduction

Dopamine release dynamics in the nucleus accumbens (NAcc) have been shown to be critical for learning the relationship between cues, actions, and outcomes ^1,2^. Across several tasks, phasic, short bursts of dopamine seem to signal errors in predicting events such as the presentation of an important cue or the delivery of reward ^1,3,4^. However, it has also been shown that *prior* to the occurrence of such events, particularly when animals are actively moving towards a desired goal, dopamine release slowly ramps up in the NAcc ^5–12^. This anticipatory dopamine ramping has been the focus of much recent work, in large part because of the rapid proposal of several alternative hypotheses for its computational role. Some propose that these ramps do not reflect a prediction error-type response, but instead signal the absolute value expectation or motivation associated with the goal ^6,6^. Alternatively, others suggest that the ramps can be explained by classical temporal difference learning algorithms ^8,9^ or that they are a correlate of the use of cognitive maps ^11^. Compounding with this controversy, there is conflicting evidence as to the origin of these anticipatory ramps. Some have argued that they are driven by matched ramps in firing in dopamine neurons ^9–11^, while others argue that they are independent of dopamine neuron spiking, and instead are generated by local circuit mechanisms in the NAcc ^6,7^, which would fit with an entirely separate computational role compared to other dopamine signals.

If these dopamine ramps are indeed generated within the NAcc, a candidate driver would be the striatal cholinergic interneurons. Previous work, including several recent mechanistic studies focusing on dopamine axon physiology, have demonstrated that acetylcholine, acting on axonal α6 nicotinic receptors, can directly drive dopamine release independently of somatic firing in midbrain neurons ^13,14^. While there are additional factors to consider about these nicotinic inputs ^15^, this would be an ideal candidate for a local circuit mechanism that could drive dopamine ramps. If this is true, then acetylcholine and dopamine signals should be positively correlated, especially during ramps, with dopamine increases lagging behind acetylcholine increases.

However, there is also a corpus of studies where cholinergic interneurons were recorded in awake behaving animals that suggest that these neurons typically pause, or “dip”, their activity when reward or reward predicting cues are presented, in opposition to dopamine^16–19^. Recent work with simultaneous striatal dopamine and acetylcholine recordings has shown that acetylcholine dips are anti-correlated during movement and reward ^20,21^. An alternative hypothesis for striatal dopamine-acetylcholine interactions which would explain these results centers on the post-synaptic response of target spiny projection neurons (SPNs), where cholinergic and dopaminergic transmission can have opposing effects on synaptic plasticity ^22,23^. According to this model, dopamine and acetylcholine dynamics should be anti-correlated, with acetylcholine dips creating a permissive window for phasic dopamine increases to drive synaptic plasticity. That said, these recording studies of cholinergic function, including the recent work with dual dopamine and acetylcholine recordings, were done in the dorsal striatum, were dopamine ramps are not typically observed ^12^. Therefore, it could still be that in NAcc there is a unique effect of acetylcholine to drive dopamine ramps.

Here, we investigated this possibility using dual fiber photometry recordings of dopamine and acetylcholine signals in the NAcc core in a simple instrumental task to assess whether these signals were positively correlated, focusing on dopamine ramps during motivated approach.

## Results

### Experimental procedures and behavioral performance

We transfected 10 male Long-Evans rats with next generation genetically-encoded sensors for dopamine and acetylcholine - rDA3m, a red-shifted dopamine sensor^24^ and gAch4h, a novel green acetylcholine sensor. These rats were implanted with optic fiber cannulas in the NAcc to allow simultaneous multi-color fiber photometry recordings of both dopamine and acetylcholine release dynamics (Figure 1A and B) ^1,20,25^. After at least 4 weeks for recovery and viral expression, rats were water restricted and started training on the behavioral task (Figure 1C).

**Figure 1.**
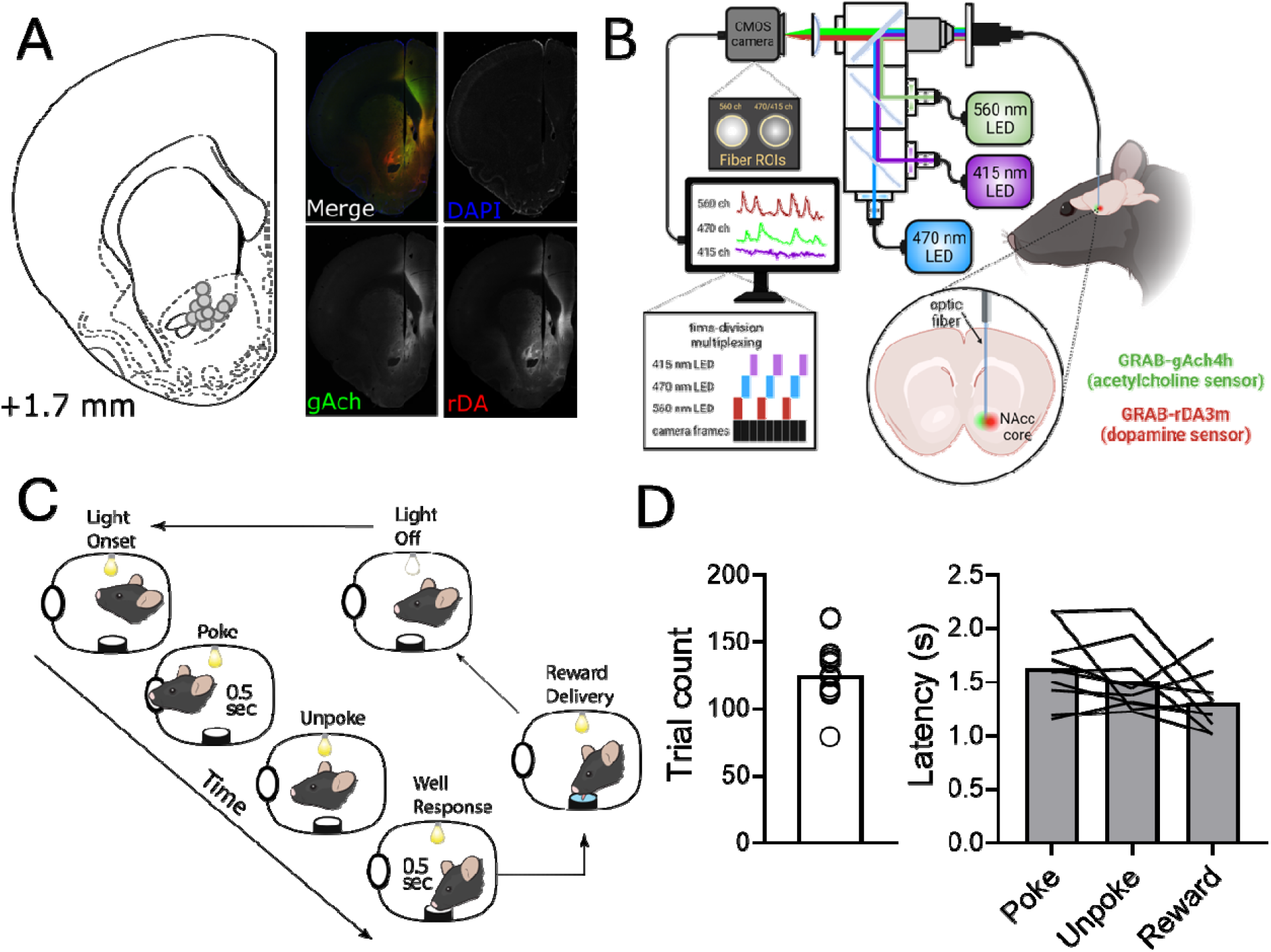
Photometry recordings, histological verification, and behavior. **A:** Location of fiber tips in the NAcc for all recorded rats (left; N=10) and representative histological microphotograph (right) with histological detection of both sensors. We would like to highlight that chicken anti-GFP antibodies were the most effective in detecting the gAch4h sensor, out of several alternatives (see Methods). **B:** Cartoon schematic of dual-color fiber photometry recording methods. **C:** Cartoon schematic of the instrumental nosepoke task. **D:** Individual and group mean responding of the rats in the behavioral task. Left panel shows the number of trials each rat performed in the one hour session, and the right panel indicates the time it took for the rats to complete each phase of the task, from light onset to nose poke (poke), from nose poke to unpoke (unpoke), and from unpoke to receiving the reward (reward).

The task was chosen to provide the simplest possible scenario in which dopamine ramps could be expected - a cued, motivated-approach behavior. On each trial, the onset of a light cue indicated that rats could perform an entry into a nose poke port, and after holding position for 0.5 seconds they could perform a second entry into a fluid well, which triggered the delivery of water rewards also after 0.5 seconds. Implanted rats quickly learned to perform this task, and we recorded acetylcholine and dopamine signals in the NAcc core during asymptotic performance (Figure 1D). All analyses reported in this study were on signals collected from one session of each rat after they reached stable performance (N=10). All sessions were limited to one hour to avoid excessive photobleaching, and the rats performed an average of 125 trials (Figure 1D). The time it took for them to executed each phase of the task was also relatively similar (Figure 1D).

The analyses reported here were conducted mainly on signals that were only detrended (to remove photobleaching artifacts), median filtered (to remove high-frequency artifacts), and z-scored (to allow for better between session and subject comparisons). We did record fluorescence elicited by 415 nm excitation, but the use of this “isosbestic” control, especially on sensors that have a shifted real isosbestic point, has recently been called into question ^26^. We found that referencing our signals to the 415 channel did not affect the interpretation of the signal dynamics (Figure S1), but we chose to continue with the most conservative approach.

### Dopamine and acetylcholine correlations vary according to task phase

Analysis of the dopamine and acetylcholine signals clearly demonstrated that they were not uniformly correlated across the different phases of the approach task (Figure 2). When we aligned the two signals to the nose poke we observed clear dopamine ramps, gradual increases in dopamine signal as the rats approached the goal, replicating several recent findings^5–7^. These ramps were significantly different from a shuffled control signal (Figure 2B), crossing the shuffled threshold well before the rats actually executed the nose poke, and their time course matched the time course of behavioral responding during this phase (Figure 1D). However, acetylcholine signals in the same period did not change, remaining statistically-similar to the shuffled control. This evidence goes against the prediction that the dopamine ramps are caused by local cholinergic depolarization of dopamine axons.

**Figure 2.**
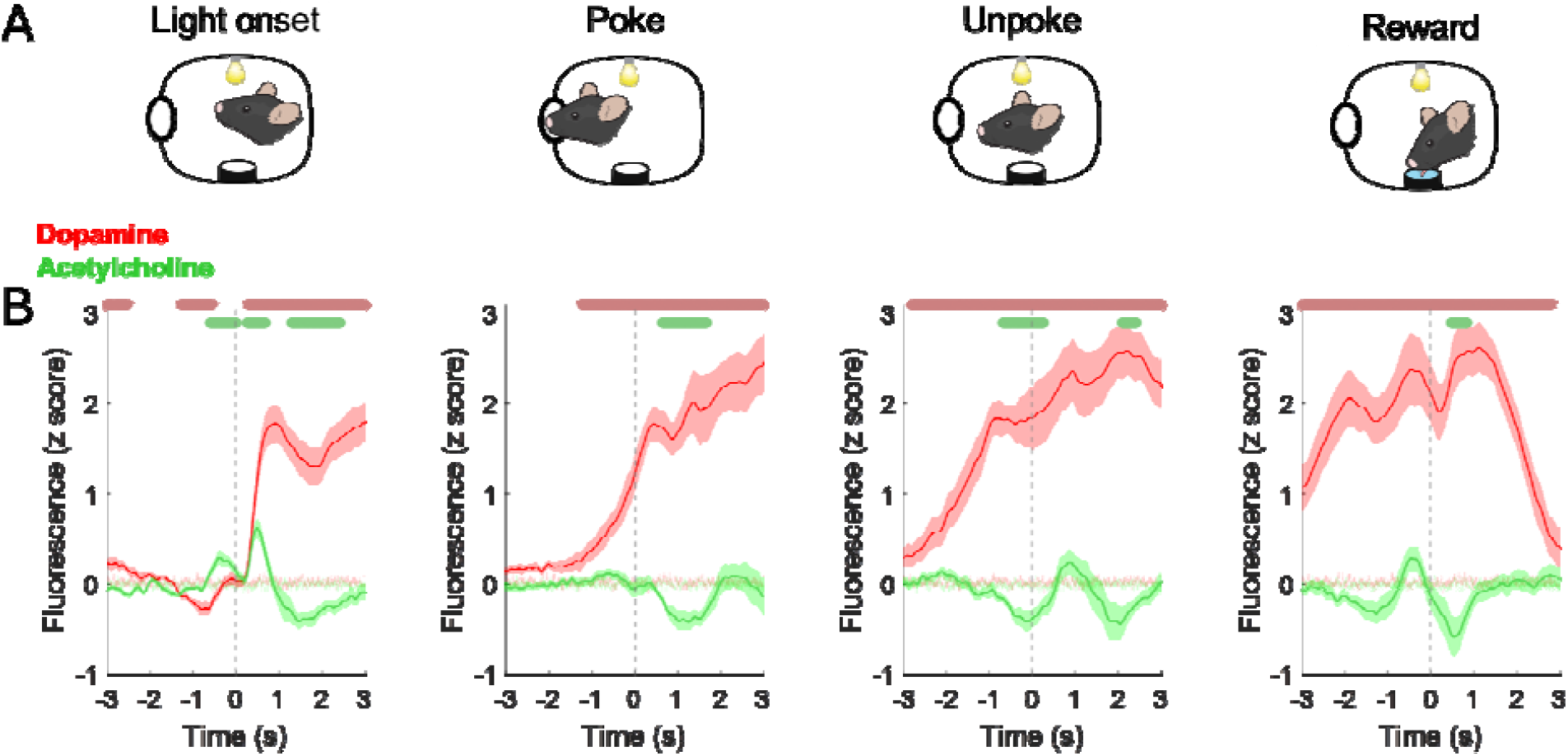
NAcc dopamine and acetylcholine dynamics during the instrumental task. **A:** Graphical representation of the task event to which each graph below is aligned (dashed gray line). **B:** Dopamine (red) and acetylcholine (green) signals aligned to the task events. Note that there is an increase in both signals immediately after light onset, a progressive dopamine ramp with no significant change in cholinergic signal prior to the nose poke, a phasic increase in dopamine right after the poke, a dip in acetylcholine centered around the unpoke and followed by an increase in dopamine, and an increase in dopamine and dip in acetylcholine immediately after the reward delivery. Data are represented as mean ± 95% CI. Light green and light red shades in the background are the SEM of the shuffled baseline control. Colored bars above graphs indicate significant difference from shuffled control using a permutation test^27^.

However, the relationship between dopamine and acetylcholine signals was very different during other task phases. When we aligned the photometry signals to the unpoke, which was the action immediately prior to reward seeking, we observed a phasic increase in dopamine and a coincidental decrease in acetylcholine (Figure 2C). The same was observed when we aligned the signals to reward port entry, with dopamine rises and acetylcholine dips occurring around the time of reward delivery (Figure 2D). This indicated that whenever the task involved a rewarded action, or reward itself, dopamine and acetylcholine signals became anticorrelated, with a characteristic burst in dopamine and dip in acetylcholine. Finally, we also found periods when the two signals were correlated. For example, when the light was turned on, indicating the start of the trial, both dopamine and acetylcholine signals showed sharp increases (Figure 2A). Although, these were also followed by a dip. Therefore, depending on the task phase and the associated behavioral processes, accumbal dopamine and acetylcholine signals can be positively correlated, negatively correlated, or uncorrelated.

### Dopamine and acetylcholine cross-correlations differ according to task phase

We next asked if the cross-correlations within and between the two signals were also different depending on task phase. This was done to rule out any potential lagged correlation that could indicate a causal relationship between the signals. We performed cross-correlation analysis on the two signals during the baseline (right before trial start), ramping (before the nosepoke), and around the light on, nosepoke, unpoke, and reward port entry timestamps, with lags computed relative to the dopamine signal. We found that during baseline, ramping, and nosepoke, the two signals had relatively weak but significant positive and negative cross-correlations, with dopamine leading the negative correlation and acetylcholine leading the positive correlation (Figure 3A). However, during the unpoke and the reward phase, signals were significantly anti-correlated across both positive and negative lags. Finally, when the trial light was turned on there was a strong positive correlation at positive lags in relation to dopamine.

**Figure 3.**
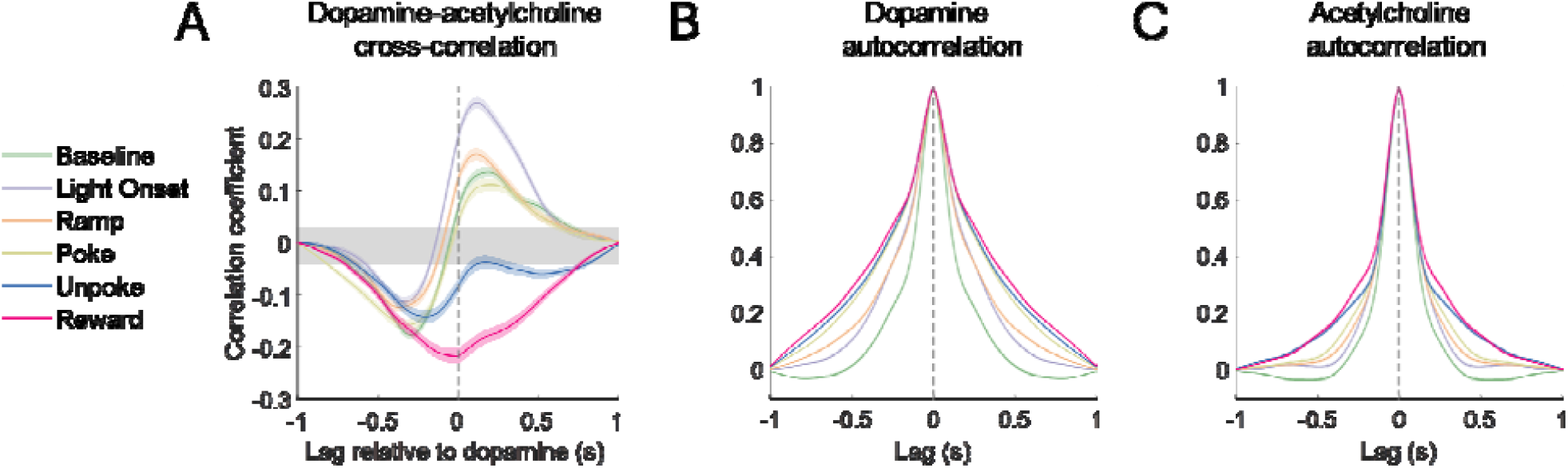
Dopamine and signals acetylcholine cross- and auto-correlations according to task phase alignment. **A:** Trial-by-trial cross-correlation between dopamine and acetylcholine signals in the NAcc during different periods of the task. Grey shade is the 95% confidence interval of the shuffled control. **B:** Autocorrelation of the dopamine signal during the same task periods. **C:** Autocorrelation of the acetylcholine signal during the same task periods. Data are represented as mean ± SEM.

We also computed the autocorrelation for each signal in the same time windows. We found that autocorrelation values for both signals also varied according to task phase, with the highest autocorrelations being observed in the task phases associated with reward (unpoke and reward) and the lowest autocorrelation being observed during baseline. There was also more task-dependent variation in autocorrelation in the dopamine signal compared to the acetylcholine signal. These analyses further confirm that the cross- and within-channel dynamics of dopamine and acetylcholine photometry differ depending on the task state.

### Dopamine and acetylcholine signal dynamics are largely independent

Finally, we wanted to explore if there was any relationship between dopamine and acetylcholine signals that could indicate a causal relationship between the two neuromodulators that spanned across task phases. For this, we removed the variance in each signal that could be explained by the variance in the other signal. In brief, we fitted a kernel function to the dopamine signal, then took the parameters of that fit and applied to acetylcholine signal, then subtracted the resulting fit from the original acetylcholine signal, and then repeated the same process with acetylcholine being the first fit and dopamine the second. The end result were dopamine and acetylcholine signals that were free of the variance explained by the dynamics of the other simultaneously recorded signal, and in which their dynamics could be compared in a scale-invariant manner (Figure 4).

**Figure 4.**
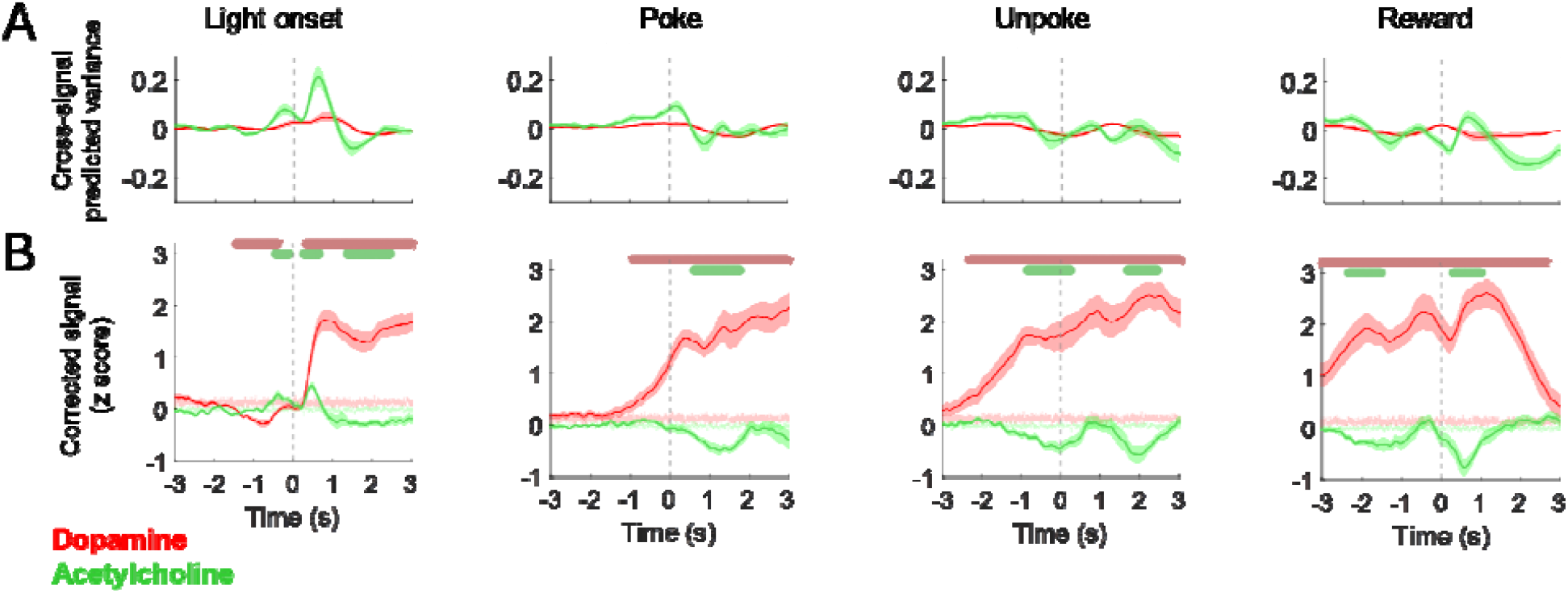
Dopamine and signals acetylcholine cross- and auto-correlations according to task phase alignment. **A:** Average traces of the acetylcholine signal variance explained by dopamine dynamics (green) and the dopamine signal variance explained by acetylcholine dynamics (red) for each task phase **B:** Dopamine (red) and acetylcholine (green) signals where the variance explained by the dynamics of the alternative signal have been removed. Note that the major patterns of activity, including anticipatory dopamine ramps and cholinergic dips during reward and rewarded action, are largely similar. Data are represented as mean ± SEM. Light green and light red shades in the background are the SEM of the shuffled baseline control. Colored bars above graphs indicate significant difference from shuffled control using a permutation test^27^.

We found that after processing the main patterns we had observed in the raw signals were all preserved. This included the dopamine ramps preceding the nosepoke, dopamine rises and acetylcholine dips during the reward-related epochs, and dopamine and acetylcholine rises to the light on. This preservation suggests that, while dopamine and acetylcholine signals may be correlated during these events, their variance is largely independent, which indicates it is unlikely that one signal directly causes changes to the other.

## Discussion

Here we simultaneously recorded dopamine and acetylcholine signals in the NAcc with genetically encoded sensors while rats performed an instrumental task that involved motivated approach. We found that dopamine and acetylcholine signal correlations vary widely depending on the task state and the behavior being executed by the rats. Essentially, dopamine and acetylcholine were positively correlated in response to the light cue that started the trial, uncorrelated during anticipatory ramps, and anticorrelated during task phases that involved reward or a directly rewarded action.

Critically, the lack of correlation between dopamine ramps and changes in acetylcholine during motivated approach demonstrates that this form of dopamine signaling is not likely driven by local acetylcholine release. Our findings contradict previous work suggesting that cholinergic interneuron activity was necessary to generate dopamine ramps ^7^. However, that study used fiber photometry to record calcium, which can be uncoupled from somatic firing and neurotransmitter release ^28^. Furthermore, the causal optogenetic evidence presented in that paper has been argued to be an artifact of direct optical stimulation of the calcium sensor ^29^. Our study, which employed a more direct measure of dopamine and acetylcholine signal in NAcc during behavior, failed to reliably find this relationship.

That said, there are methodological limits to consider when interpreting our findings. Photometry recordings sample a relatively large area of neural tissue, with no cell-type specificity, so there is still the possibility that there may be variations in dopamine-acetylcholine interactions at the cellular and subcellular levels that were not captured with our methods. Nevertheless, our findings are well in line with most of the previous literature, and signals recorded at the level of photometry are typically highly correlated and causally linked to behavioral performance^3,9,26^.

Our findings are also in line with previous electrophysiological and photometry studies of cholinergic transmission in the dorsal striatum. Cholinergic interneurons in the dorsal striatum tend to pause during the presentation of reward and reward-predicting cues, while dopamine neurons tend to burst in the same conditions ^16^, and dopamine and acetylcholine release in the dorsal striatum is anti-correlated during reward^20,21^, both patterns that fit with our dual photometry results in the NAcc. Additionally, individual striatal cholinergic interneurons have also been found to burst, or burst and then dip, in response to cued events, similarly to what we observed in response to the light onset^16–19^. This suggests that the general local circuit structure governing dopamine and acetylcholine release in the NAcc is also somewhat similar to what has been described for the dorsal striatum.

If acetylcholine changes are not a prerequisite for dopamine ramping, then this suggests that, pre-synaptically, dopamine ramps likely share the same mechanisms as other dopamine signaling events, like classical reward prediction errors. However, it is conspicuous that we observed an anti-correlation between dopamine and acetylcholine in the precise epochs that dopamine should be signaling reward prediction errors and, presumably, driving reward-related learning. Specifically, cholinergic dips and dopamine increases coincide with events that are intrinsically or have been previously directly associated with value. This may indicate that, even if the ramps and prediction error signals are both generated by the same presynaptic mechanisms, they may engage different postsynaptic targets.

For example, the anticorrelation pattern fits well with the finding that dopamine and acetylcholine exert opposing effects on each major classical SPN pathway. In the direct pathway dopamine acts on D1 receptors while acetylcholine acts on M4 receptors, respectively boosting and decreasing synaptic plasticity^22^. Conversely, in the indirect pathway, dopamine acts on D2 receptors and acetylcholine acts on M1 receptors, which also exert opposing effects on plasticity in these SPNs ^23^. It has been proposed that this oppositional relationship creates a tripartite condition for synaptic plasticity to occur in each SPN pathway, where learning is set to occur primarily when there is a coincidental dopamine burst, acetylcholine dip, and post-synaptic depolarization^30^. That said, the real situation is almost certainly more complex than this, as both modulators also act on different interneurons and on dopamine axons themselves^14,15,31,32^, and there is compounding evidence that both direct and indirect SPNs are dynamically co-active during learning and decision-making^33–36^. The mechanistic model described previously is intended as an initial heuristic for investigating dopamine and acetylcholine interactions in subsequent studies.

Within this framework, the fact that dopamine and acetylcholine are *not* anticorrelated during motivated approach and salient cue exposure is very interesting. This suggests that during these epochs the combined post-synaptic effect of both neuromodulators may be quite different. It is also an indication that dopamine ramps and cue responses are indeed mechanistically different from classical reward prediction error responses, at least in terms of how they modulate target cells. While all these dopamine responses can be conceptualized as prediction errors, reward-based or otherwise ^1,9^, they clearly drive different behaviors, and therefore it would make sense that they engage different post-synaptic cellular mechanisms. Regarding specifically the dip in acetylcholine during dopamine reward prediction error signaling, it could be that the dips reflect the associative salience of the actions and reward and creates a critical window where dopamine can drive associative learning-related plasticity. This possibility should be actively explored in future work.

The highly correlated responses to the light cue are harder to interpret within the confines of our task. This cue is related to reward availability and also indicates that the rat can initiate an action, therefore the cholinergic responses could be related to both to an action sequence initiation and value. However, it is worth noting that the light onset differs from other elements of the task as being the only highly salient cue that is outside of the rat’s control, and thus the dopaminergic and cholinergic responses to this event may be dominated by physical salience or a sensory prediction error. Future work with more complex tasks will be needed to disambiguate the nature of these responses.

In conclusion, our findings indicate that the correlation between dopamine and acetylcholine release in the NAcc is heavily dependent on the precise timing and type of behavioral processes, even in relatively simple tasks. Dopamine increases in response to most events in this task, but acetylcholine dips during events directly related to reward and peaks during salient trial-setting cues. Importantly, anticipatory dopamine ramps are not coincidental with major changes in cholinergic signals. This pattern of results suggests that different behavior-related dopamine signals may induce specific post-synaptic effects in NAcc neurons depending on their interaction with acetylcholine dynamics.

## Methods

### Materials and correspondence

All data and code displayed in this manuscript will be made available upon request. Additional information on materials and protocols are available upon request to Geoffrey Schoenbaum.

### Experimental Model and Subject Details

Experiments were performed on a total of 10 male Long-Evans rats (>3 months of age at the start of the experiment, Charles River Laboratories) housed on a 12 hr light/dark cycle at 25 °C. Rats were water restricted (10 minutes/day) for the duration of the experiments and were tested at the NIDA-IRP in accordance with NIH guidelines determined by the Animal Care and Use Committee, which approved all procedures. All rats had *ad libitum* access to rat chow in their home cages for the duration of the experiments. Behavior was performed during the light phase of the light/dark schedule. The lack of female rats, due to logistical issues and the fact that males performed better with the head implants, is a potential limitation of this study.

### Surgical procedures

Rats were anesthetized with 1-2% isoflurane and prepared for aseptic surgery. They received unilateral infusions of AAV2/9-hSyn-rDA3m and AAV2/9-hSyn-gAch4h into the NAcc (AP +1.7 mm, ML + or -1.7 mm, and DV -6.3 and -6.2 mm from the brain surface). Viruses were mixed in a small tube and a total 0.7 μL of this mixture was delivered in each site at 0.1 μL/min via an infusion pump. Optic fiber cannulas (200 μm diameter; Neurophotometrics, CA) were implanted in each site in the location of the second (most dorsal) viral infusion. All viruses were obtained from BrainVTA. Exposed fiber ferrules and a protective black 3D-printed headcap were secured to the skull with dental cement. After surgery, rats were given Cephalexin (15 mg/kg po qd) for two weeks to prevent any infection.

### Dual color fiber photometry

Fluorescent dopamine and acetylcholine signals were recorded using dual-color fiber photometry. General methods were similar to what was described previously^1^. In brief, custom fiber optic patch cables (200 μm diameter, 0.37 NA, Doric Lenses, Canada) were attached to the optic fiber ferrules on the rats with brass sleeves (Thorlabs, NJ). Fibers were shielded and secured with a custom 3D-printed headcap-swivel shielding. Recordings were conducted using an FP3002 system (Neurophotometrics, CA), by providing 560 (active green signal), 470 (active green signal) and 415 nm (isosbestic reference) excitation light through the patch cord in interleaved LED pulses at 150 Hz (50 Hz acquisition rate for each channel). The light was reflected through a dichroic mirror and onto a 20× Olympus objective. Excitation power was measured at ∼70-90 μW at the tip of the patch cord. Emitted fluorescent light was captured via a high quantum efficiency CMOS camera. Signals were acquired and synchronized with behavioral events using Bonsai^37^.

Signals were processed using custom scripts in Python and MATLAB (MathWorks, MA). We filtered raw fluorescence signals from each of the 470 nm(active), 560 nm (active), and 415 nm (reference) channels with a causal median filter and a second-order Butterworth low-pass filter with a cutoff frequency of 5 Hz. Next, each channel data was fitted with a double exponential function, and the fitted data was subtracted from the original signal which removed the exponential decay artifact caused by photobleaching. The resulting signal was z-scored for each trial, using the three seconds before each trial onset as a baseline. For the supplemental reference control analysis, the reference (415 nm) channel data was fitted to each active signal using second-order polynomial regressions, and the fitted data was subsequently subtracted from the active channel and divided by the exponential fit of the active channel.

### Signal analyses

Cross- and autocorrelations were conducted on one second windows using MATLAB’s *x-corr* function. Periods for the execution of the analyses started at ∼2 seconds before light onset (baseline), immediately after light onset (light on), one second preceding nose poke (ramp), immediately after nose poke (poke), 0.5 second before unpoke (unpoke), and immediately after reward delivery (reward). The 95% confidence interval was derived by repeatedly calculating Pearson’s r after one of the photometry signals was shifted in time (aligned to the light onset and spanning the whole trial) and then extracting the 2.5th and 97.5th percentiles across the correlation window for each bin, similar to what has been used previously^20^.To address whether the dynamics of ACh and DA to each event derive from the other signaling, we isolated the component of one signal that could not be predicted by the other signal by regressing the data of one neurotransmitter to predict the other and subtracting this predicted component from the original signal.

To address whether the dynamics of dopamine and acetylcholine influence each other, we isolated the component of one signal that could not be predicted by the other signal by regressing the data of one neurotransmitter to predict the other and subtracting this predicted component from the original signal. The regression was done by using the data, *x*, in the past 2 seconds to predict the current response of the other neurotransmitter, *y*, using a double exponential kernel:

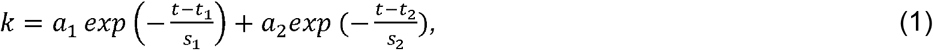

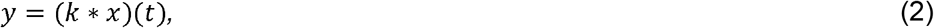

where (*k* * *x*)(*t*) indicates the linear convolution between data *x* and kernel *k*. Parameters *a*_1_ and *a*_2_ control the amplitude, *t*_1_ *a*nd *t*_2_ represent time shifts for each phase, and time constants *s*_1_ and *s*_2_ govern the sharpness.

We optimized these parameters for each session by minimizing mean squared error. With the optimized parameters, we were able to predict one signal based on the historical data of the other through convolution with the fitted kernel. Subsequently, this predicted component was removed from the original signal and tested to see if the response to each event was changed afterward.

### Behavioral apparatus and instrumental nose poke task

Rats were trained and tested at least four weeks after the surgeries. Water was restricted to ∼10 min free access every day for at least two days prior to training initiation. During training, they received their water ration after their daily session. All behavior experiments were conducted in custom-built aluminum chambers approximately 18’ on each side with sloping walls narrowing to an area of 12’ x 12’ at the bottom. A central nose poke port consisting of a small hemicylinder accessible was located about 2 cm above a fluid well, and higher up on the same wall were mounted two lights. Trial availability was signaled by the illumination of the panel lights. When these lights were on, if rats performed a 500 ms nosepoke into the odor port and then made a response into the fluid well and hold for 500 ms, they would receive a ∼0.05 mL drop of water. Rats were trained until they could reliably perform over 75 trials in a one hour period.

### Histological procedures

After completion of the experiment, rats were perfused with chilled phosphate buffer saline (PBS) followed by 4% paraformaldehyde in PBS. The brains were post-fixed in 4% PFA for at least 24 hours then immersed in 30% sucrose in PBS until they sank, and then frozen. The brains were sliced at 50 μm, stained with DAPI (Vectashield-DAPI, Vector Lab, Burlingame, CA), and processed for immunohistochemical detection of green and red fluorescent proteins (Figures 1A and S4B and C). For immunohistochemistry, the brain slices were first washed with PBS (5x10 mins), blocked in 4% BSA with 0.3% Triton X-100 in PBS, and then incubated with anti-GFP (1/1000, RT, overnight, chicken anti-GFP, ab13970, Abcam USA, Waltham, MA) and anti-RFP antibodies (1/1000, RT, overnight, rabbit anti-DsRed, 632496, Takara Bio USA, Madison, WI), followed by Alexa-488 (1/100, RT, 2h, Donkey anti-chicken Alexa Fluor 488, ab2340375, Abcam, Waltham, MA) and Alexa-594 (1/100, RT, 2h, Donkey anti-rabbit Alexa Fluor 594, ab2340621, Abcam, Waltham, MA) secondary antibodies. We want to call attention that the chicken anti-GFP antibody used here was the most successful at detecting the gAch4.0h sensor. We tested several alternatives (data not shown), made in different species and from different vendors, and highly recommend the use of this antibody for this sensor. Fluorescent microscopy images of the slides were acquired with an Olympus VS120 microscope (Figure 1A).

### Statistical analyses

Statistical analyses were performed in MATLAB. Significant differences between the signals and shuffled controls were conducted using permutation tests ^27^, with a consecutive threshold of fifteen, ten thousand permutations, and statistical significance set at *P*<0.05.

## Acknowledgements

This work was supported by the Intramural Research Program at the National Institute on Drug Abuse (Z1A DA000587 to GS) and by the NIH BRAIN Initiative (NINDS U01NS120824 to Y.L.). We thank Shiliang Steven Zhang and the NIDA IRP Histology and Imaging Core for assistance with antibody testing and histological processing. The opinions expressed in this article are the authors’ own and do not reflect the view of the NIH/DHHS.

## Author contributions

KMC, ZZ, and GS designed experiments, interpreted results, and wrote the manuscript. YZ, GL, and YL provided important and unpublished reagents that were essential for the photometry experiments. KMC and ZZ performed the photometry experiments and analyzed the data. KMC wrote the first draft of the manuscript. GS supervised the research.

## Supplemental information

**Figure S1.**
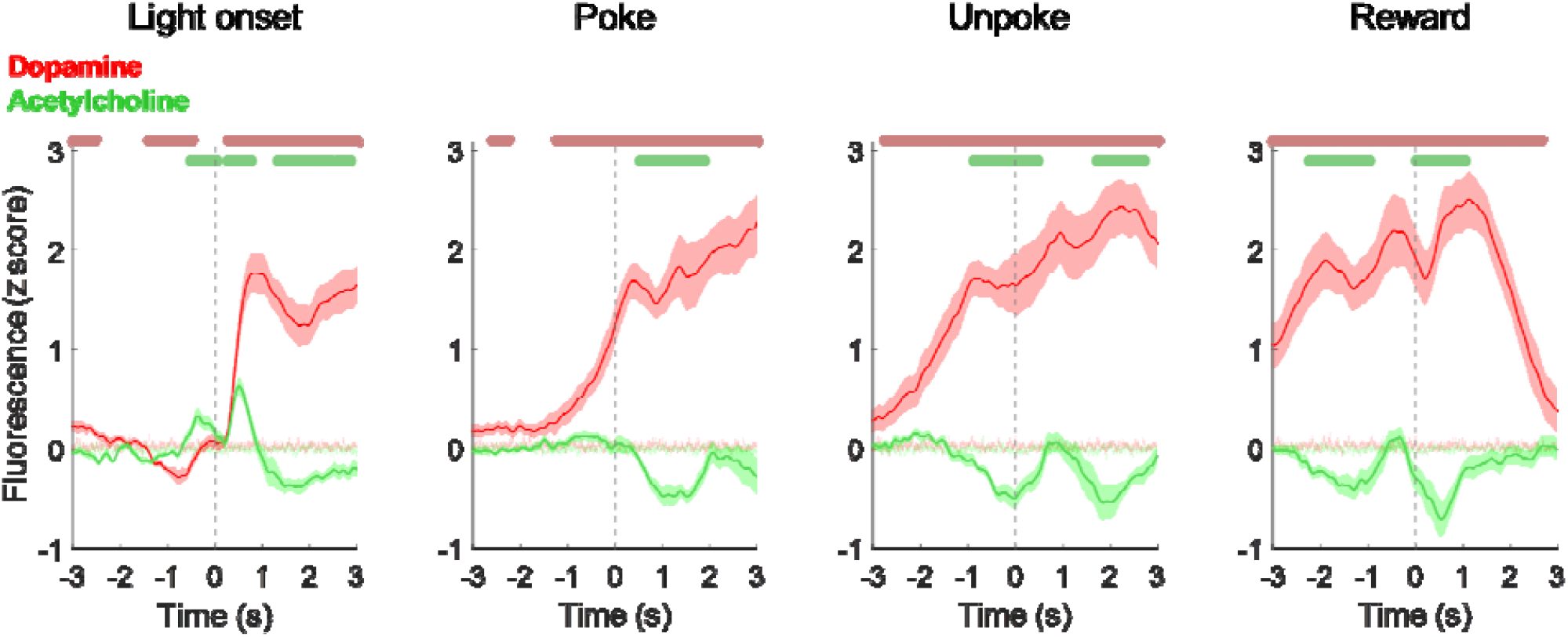
Referencing signals with 415 fluorescence does not change their dynamics. The same graphs as presented in Figure 1 but with referencing to the 415 channel with dopamine in red and acetylcholine in green. Note that the signal patterns do not significantly change with the referencing process. Data are represented as mean ± SEM. Light green and light red shades in the background are the SEM of the shuffled baseline control. Colored bars above graphs indicate significant difference from shuffled control using a permutation test^27^.

## References

1. Costa, K.M., Raheja, N., Mirani, J., Sercander, C., and Schoenbaum, G. (2023). Striatal dopamine release reflects a domain-general prediction error. bioRxiv, 2023.08.19.553959. 10.1101/2023.08.19.553959.

2. Day, J.J., Roitman, M.F., Wightman, R.M., and Carelli, R.M. (2007). Associative learning mediates dynamic shifts in dopamine signaling in the nucleus accumbens. Nat Neurosci 10, 1020–1028. 10.1038/nn1923.

3. Menegas, W., Babayan, B.M., Uchida, N., and Watabe-Uchida, M. (2017). Opposite initialization to novel cues in dopamine signaling in ventral and posterior striatum in mice. Elife 6, e21886. 10.7554/eLife.21886.

4. Costa, K.M., and Schoenbaum, G. (2022). Dopamine. Current Biology 32, R817–R824. 10.1016/J.CUB.2022.06.060.

5. Howe, M.W., Tierney, P.L., Sandberg, S.G., Phillips, P.E.M., and Graybiel, A.M. (2013). Prolonged dopamine signalling in striatum signals proximity and value of distant rewards. Nature 500, 575–579. 10.1038/nature12475.

6. Mohebi, A., Pettibone, J.R., Hamid, A.A., Wong, J.-M.T., Vinson, L.T., Patriarchi, T., Tian, L., Kennedy, R.T., and Berke, J.D. (2019). Dissociable dopamine dynamics for learning and motivation. Nature 570, 65–70. 10.1038/s41586-019-1235-y.

7. Mohebi, A., Collins, V.L., and Berke, J.D. (2023). Accumbens cholinergic interneurons dynamically promote dopamine release and enable motivation. eLife 12, e85011. 10.7554/eLife.85011.

8. Farrell, K., Lak, A., and Saleem, A.B. (2022). Midbrain dopamine neurons signal phasic and ramping reward prediction error during goal-directed navigation. Cell Rep 41, 111470. 10.1016/j.celrep.2022.111470.

9. Kim, H.R., Malik, A.N., Mikhael, J.G., Bech, P., Tsutsui-Kimura, I., Sun, F., Zhang, Y., Li, Y., Watabe-Uchida, M., Gershman, S.J., et al. (2020). A Unified Framework for Dopamine Signals across Timescales. Cell 183, 1600-1616.e25. 10.1016/j.cell.2020.11.013.

10. de Jong, J.W., Liang, Y., Verharen, J.P.H., Fraser, K.M., and Lammel, S. (2024). State and rate-of-change encoding in parallel mesoaccumbal dopamine pathways. Nat Neurosci 27, 309–318. 10.1038/s41593-023-01547-6.

11. Guru, A., Seo, C., Post, R.J., Kullakanda, D.S., Schaffer, J.A., and Warden, M.R. (2020). Ramping activity in midbrain dopamine neurons signifies the use of a cognitive map. Preprint at bioRxiv, 10.1101/2020.05.21.108886 10.1101/2020.05.21.108886.

12. Chow, J.J., Pitts, K.M., Schoenbaum, A., Costa, K.M., Schoenbaum, G., and Shaham, Y. (2024). Different Effects of Peer Sex on Operant Responding for Social Interaction and Striatal Dopamine Activity. J Neurosci 44, e1887232024. 10.1523/JNEUROSCI.1887-23.2024.

13. Rice, M.E., and Cragg, S.J. (2004). Nicotine amplifies reward-related dopamine signals in striatum. Nat Neurosci 7, 583–584. 10.1038/nn1244.

14. Kramer, P.F., Brill-Weil, S.G., Cummins, A.C., Zhang, R., Camacho-Hernandez, G.A., Newman, A.H., Eldridge, M.A.G., Averbeck, B.B., and Khaliq, Z.M. (2022). Synaptic-like axo-axonal transmission from striatal cholinergic interneurons onto dopaminergic fibers. Neuron 110, 2949-2960.e4. 10.1016/j.neuron.2022.07.011.

15. Zhang, Y.-F., Luan, P., Qiao, Q., He, Y., Zatka-Haas, P., Zhang, G., Lin, M.Z., Lak, A., Jing, M., Mann, E.O., et al. (2024). An axonal brake on striatal dopamine output by cholinergic interneurons. Preprint at bioRxiv, 10.1101/2024.02.17.580796 10.1101/2024.02.17.580796.

16. Morris, G., Arkadir, D., Nevet, A., Vaadia, E., and Bergman, H. (2004). Coincident but Distinct Messages of Midbrain Dopamine and Striatal Tonically Active Neurons. Neuron 43, 133–143. 10.1016/j.neuron.2004.06.012.

17. Apicella, P., Scarnati, E., and Schultz, W. (1991). Tonically discharging neurons of monkey striatum respond to preparatory and rewarding stimuli. Exp Brain Res 84, 672–675. 10.1007/BF00230981.

18. Deffains, M., and Bergman, H. (2015). Striatal cholinergic interneurons and cortico-striatal synaptic plasticity in health and disease. Movement Disorders 30, 1014–1025. 10.1002/mds.26300.

19. Aosaki, T., Kimura, M., and Graybiel, A.M. (1995). Temporal and spatial characteristics of tonically active neurons of the primate’s striatum. Journal of Neurophysiology. 10.1152/jn.1995.73.3.1234.

20. Krok, A.C., Maltese, M., Mistry, P., Miao, X., Li, Y., and Tritsch, N.X. (2023). Intrinsic dopamine and acetylcholine dynamics in the striatum of mice. Nature 621, 543–549. 10.1038/s41586-023-05995-9.

21. Chantranupong, L., Beron, C.C., Zimmer, J.A., Wen, M.J., Wang, W., and Sabatini, B.L. (2023). Dopamine and glutamate regulate striatal acetylcholine in decision-making. Nature 621, 577–585. 10.1038/s41586-023-06492-9.

22. Shen, W., Plotkin, J.L., Francardo, V., Ko, W.K.D., Xie, Z., Li, Q., Fieblinger, T., Wess, J., Neubig, R.R., Lindsley, C.W., et al. (2015). M4 Muscarinic Receptor Signaling Ameliorates Striatal Plasticity Deficits in Models of L-DOPA-Induced Dyskinesia. Neuron 88, 762–773. 10.1016/j.neuron.2015.10.039.

23. Shen, W., Tian, X., Day, M., Ulrich, S., Tkatch, T., Nathanson, N.M., and Surmeier, D.J. (2007). Cholinergic modulation of Kir2 channels selectively elevates dendritic excitability in striatopallidal neurons. Nat Neurosci 10, 1458–1466. 10.1038/nn1972.

24. Zhuo, Y., Luo, B., Yi, X., Dong, H., Miao, X., Wan, J., Williams, J.T., Campbell, M.G., Cai, R., Qian, T., et al. (2023). Improved green and red GRAB sensors for monitoring dopaminergic activity in vivo. Nat Methods. 10.1038/s41592-023-02100-w.

25. Martianova, E., Aronson, S., and Proulx, C.D. (2019). Multi-Fiber Photometry to Record Neural Activity in Freely-Moving Animals. J Vis Exp. 10.3791/60278.

26. Simpson, E.H., Akam, T., Patriarchi, T., Blanco-Pozo, M., Burgeno, L.M., Mohebi, A., Cragg, S.J., and Walton, M.E. (2024). Lights, fiber, action! A primer on in vivo fiber photometry. Neuron 112, 718–739. 10.1016/j.neuron.2023.11.016.

27. Jean-Richard-dit-Bressel, P., Clifford, C.W.G., and McNally, G.P. (2020). Analyzing Event-Related Transients: Confidence Intervals, Permutation Tests, and Consecutive Thresholds. Front. Mol. Neurosci. 13. 10.3389/fnmol.2020.00014.

28. Legaria, A.A., Matikainen-Ankney, B.A., Yang, B., Ahanonu, B., Licholai, J.A., Parker, J.G., and Kravitz, A.V. (2022). Fiber photometry in striatum reflects primarily nonsomatic changes in calcium. Nat Neurosci 25, 1124–1128. 10.1038/s41593-022-01152-z.

29. Taniguchi, J., Melani, R., Chantranupong, L., Wen, M.J., Mohebi, A., Berke, J., Sabatini, B., and Tritsch, N. (2024). Comment on ‘Accumbens cholinergic interneurons dynamically promote dopamine release and enable motivation.’ Preprint at bioRxiv, 10.1101/2023.12.27.573485 10.1101/2023.12.27.573485.

30. Reynolds, J.N.J., Avvisati, R., Dodson, P.D., Fisher, S.D., Oswald, M.J., Wickens, J.R., and Zhang, Y.-F. (2022). Coincidence of cholinergic pauses, dopaminergic activation and depolarisation of spiny projection neurons drives synaptic plasticity in the striatum. Nat Commun 13, 1296. 10.1038/s41467-022-28950-0.

31. Clarke, R., and Adermark, L. (2015). Dopaminergic regulation of striatal interneurons in reward and addiction: Focus on alcohol. Neural Plasticity 2015. 10.1155/2015/814567.

32. Ford, C.P. (2014). The role of D2-autoreceptors in regulating dopamine neuron activity and transmission. Neuroscience 282, 13–22. 10.1016/j.neuroscience.2014.01.025.

33. Cui, G., Jun, S.B., Jin, X., Pham, M.D., Vogel, S.S., Lovinger, D.M., and Costa, R.M. (2014). Concurrent activation of striatal direct and indirect pathways during action initiation. Nature 494, 238–242. 10.1038/nature11846.

34. Soares-Cunha, C., Coimbra, B., Sousa, N., and Rodrigues, A.J. (2016). Reappraising striatal D1-and D2-neurons in reward and aversion. Neuroscience & Biobehavioral Reviews 68, 370–386. 10.1016/j.neubiorev.2016.05.021.

35. Deseyve, C., Domingues, A.V., Carvalho, T.T.A., Armada, G., Correia, R., Vieitas-Gaspar, N., Wezik, M., Pinto, L., Sousa, N., Coimbra, B., et al. (2024). Nucleus accumbens neurons dynamically respond to appetitive and aversive associative learning. Journal of Neurochemistry 168, 312–327. 10.1111/jnc.16063.

36. Zachry, J.E., Kutlu, M.G., Yoon, H.J., Leonard, M.Z., Chevée, M., Patel, D.D., Gaidici, A., Kondev, V., Thibeault, K.C., Bethi, R., et al. (2024). D1 and D2 medium spiny neurons in the nucleus accumbens core have distinct and valence-independent roles in learning. Neuron 112, 835-849.e7. 10.1016/j.neuron.2023.11.023.

37. Lopes, G., Bonacchi, N., Frazão, J., Neto, J.P., Atallah, B.V., Soares, S., Moreira, L., Matias, S., Itskov, P.M., Correia, P.A., et al. (2015). Bonsai: an event-based framework for processing and controlling data streams. Frontiers in neuroinformatics 9. 10.3389/FNINF.2015.00007.

